# Bioresorbable, Injectable In Situ-Formed Electroactive mRNA-Delivering Device for Precision Gene Therapy after Myocardial Infarction

**DOI:** 10.64898/2026.10.29.758467

**Authors:** Umut Aydemir, Sevilay Sahoglu Göktas, Upala Dasmahapatra, Amit Singh Yadav, Karin Hellman, Sumit Sumit, Fredrik Ek, Anders Arner, Anja Meissner, Martin Hjort, Roger Olsson

## Abstract

Myocardial infarction (MI) causes irreversible cardiomyocyte loss and triggers inflammatory and fibrotic remodeling that promote the progression to heart failure. Although regenerative gene therapy offers a promising route to cardiac repair, clinical translation has been limited because of the lack of targeted, minimally invasive, and clinically scalable delivery strategies. Here we report an injectable, bioresorbable in situ-formed electroactive device that enables local mRNA delivery through the pericardial space, a clinically established route of access. Following administration, the formulation self-assembles into a transient conductive mRNA-carrying drape electrode over the injured heart, enabling in vivo electrotransfection, and is then bioresorbed, eliminating the need for retrieval. In animal models, the formulation was well tolerated, did not disrupt cardiac contractile activity, and enabled targeted mRNA transfer. In a mouse model of MI, localized delivery of vascular endothelial growth factor (VEGF-A) mRNA attenuated adverse remodeling, improved cardiac function (e.g., stroke volume and ejection fraction were fully recovered), and established a minimally invasive strategy for localized regenerative gene delivery to the injured heart.

## Introduction

Cardiovascular disease remains the leading cause of mortality worldwide, and myocardial infarction (MI) continues to be a major driver of heart failure despite substantial advances in reperfusion strategies, pharmacological therapies, and device-based care. A central reason for this persistent clinical burden is the limited regenerative capacity of the adult mammalian heart^1^. The damaged myocardium is gradually replaced by fibrotic scar tissue, and in advanced heart failure, transplantation remains the only definitive treatment, but its impact is constrained by donor-organ scarcity, lifelong immunosuppression, and procedural complexity.

Gene therapy offers a particularly attractive route to cardiac regeneration because it can induce local expression of therapeutic proteins directly within injured myocardium^2^. Among candidate factors, angiogenic and reparative molecules such as VEGF-A have shown considerable promise^3^. However, the principal barrier to clinical translation is no longer simply the identification of therapeutic genes. Rather, it is the development of delivery strategies that are safe, efficient, spatially targeted, compatible with the post-infarction therapeutic window, and scalable for clinical use^4^.

In the EPICCURE trial^5^, VEGF-A mRNA (AZD8601) was delivered, demonstrated the clinical feasibility of therapeutic mRNA delivery for cardiac repair, but administration required open-chest access and thirty epicardial injections during cardioplegic arrest. Lipid nanoparticles and viral vectors are leading platforms for nucleic acid delivery; however, their therapeutic application, remains limited by toxicity, immune activation, and off-target biodistribution^6^.

Preclinical studies in swine^7^ and rats^8^ support cardiac gene electrotransfer as a promising approach for localized myocardial delivery. However, existing cardiac electroporation methods still depend on direct cardiac access, largely because current electrode technologies require open-chest exposure for effective application. Thus, although these approaches establish feasibility, they also underscore invasiveness as a major obstacle for translation^9^.

To address this limitation, we developed an injectable, in situ-forming, electroactive mRNA- carrying device designed to form a conformal intrapericardial drape for localized cardiac gene delivery. The pericardial cavity provides an attractive access route because intrapericardial delivery is clinically established in humans and can be performed minimally invasively under ultrasound, fluoroscopic, or wire guidance^10^. Moreover, intrapericardial delivery of biomaterials has shown good tolerability in clinically relevant porcine models^11^. We reasoned that combining this access route with mild, localized point-source electroporation in a monopolar configuration ^12^could enable spatially targeted mRNA delivery to the epicardial surface and underlying myocardium, while avoiding open-chest electrode placement. In doing so, this strategy addresses two major barriers that have limited the broader clinical translation of electroporation: the difficulty of achieving localized electric-field delivery in deep tissues in concert with mRNA delivery and the procedural invasiveness required to access tissues, despite the widespread in vitro use of electroporation for efficient mRNA delivery.^9^

Our approach builds on our recently reported bioresorbable injectable cardiac stimulator, BICS, a nanoparticle formulation that self-assembles within the pericardial cavity into a conductive hydrogel capable of modulating cardiac activity^13^. BICS adheres to the heart, exhibits a Young’s modulus of 21 kPa, does not interfere with normal cardiac function, and was fully bioresorbed within one week without detectable toxicity. These properties suggested that the BICS nanoparticle matrix could serve in a point-source electroporation^12,14^ not only as an electroactive interface but also as a transient depot for therapeutic cargo. Specifically, we asked whether the matrix, composed of negatively charged A5 and zwitterionic ETE-BuSA, could accommodate negatively charged nucleic acids such as mRNA. If compatible with in vivo self-assembly, such cargo could be retained within the forming drape, released through charge-mediated interactions, and delivered to nearby cells via mild, targeted electroporation.

Here, we loaded mRNA into the nanoparticle dispersion, injected it, and used the resulting electroactive, in situ-formed drape to generate localized electrical pulses, thereby enabling precise and mild gene electrotransfer following minimally invasive intrapericardial administration in vivo. Proof-of-concept studies in ex vivo chicken and porcine hearts and in vivo in zebrafish hearts showed efficient electrotransfection of GFP- and mCherry-encoding mRNAs, with precise targeting and excellent tolerability in zebrafish despite their small heart size. We then assessed therapeutic efficacy and tolerability in a mouse model of MI, using VEGF-A-encoding mRNA. By the study endpoint, ejection fraction and stroke volume had recovered to within the normal physiological range observed in healthy mice.

More broadly, this platform provides a framework for localized, transient, and bioresorbable delivery of nucleic acids to the heart, with potential relevance to regenerative therapy after MI.

## Materials & Methods

### Preparation of A5:ETE-BuSA mRNA injectable solution

PEDOT-S (A5) and ETE-BuSA were synthesized as previously described^13,15^. The ETE-BuSA trimer was dissolved in Milli-Q water at a concentration of 40 mg mL^-1^. To this solution, PEDOT- S (A5) was added, resulting in an A5:ETE-BuSA 20:40 mg mL^-1^ solution. For electrotransfection studies, the A5:ETE-BuSA solution was mixed 1:1 (v/v) with 1.0 mg mL^-1^ mRNA (GFP-mRNA (Ribopro, Batch #001763, RIB-01672), mCherry mRNA, or VEGF-A mRNA (Messenger Bio). The formulation was prepared immediately before injection to preserve mRNA integrity.

### General gene electrotransfer procedure

Following injection of the polymer dispersion into the target agarose/tissue, the resulting A5:ETE- BuSA scaffold was connected externally to an Au-coated tungsten electrode and used as the point- source electrode (anode). Before pulse delivery, ETE-BuSA was volumetrically electrofunctionalized onto the self-assembled A5 scaffold by applying 1 V DC for 5 mins against an Ag/AgCl return electrode using a Keithley 2612B SourceMeter (Keithley Instruments), positioned at a distance. After approximately one minute, square-wave voltage pulses were then delivered using the parameters indicated in the text, figure, or figure legends: 60 or 90 V, with pulse durations of 200 µs, 2 ms, 20 ms, or 200 ms, at 1 Hz for 20 s, using a Grass stimulator (Astro- Med). Reversing the electrode polarity makes the cathode the point-source electrode (Nepagene).

### In vitro mediated gene electrotransfer

U87 cells were cultured in T225 flasks until reaching 90 % confluence in Dulbecco’s modified Eagle’s medium (DMEM) (Gibco) supplemented with 10% fetal bovine serum (FBS) (Gibco), 100 U/mL penicillin, 100 µg mL^-1^ streptomycin, and 1 % non-essential amino acids at 37 °C with 5 % CO₂. Prior to use, cells were trypsinized and resuspended in melted agarose (0.6% in complete DMEM without phenol red) at 37°C. The agarose–cell mixture (1 × 10⁶ cells mL^-1^) was cast into pre-fabricated 1 cm × 1 cm molds. After solidification, 2 µL of A5:ETE-BuSA formulated with GFP mRNA was injected laterally into the agarose blocks using a Hamilton syringe. For dose– response experiments, the relative mRNA concentration was varied to 0.5, 1.0, and 1.5 µg/mL.

Following injection of A5:ETE-BuSA + mRNA and electrofunctionalization, electrotransfection was subsequently performed using square voltage pulses at 60 V, 1 Hz, and a 2 ms pulse width for 20 seconds. After electrotransfection, agarose blocks were transferred to CO₂-independent medium in 6-well plates and incubated overnight at 37.5°C. Blocks were then fixed overnight in 4% paraformaldehyde, rinsed in PBS, cryoprotected in 25 % sucrose in PBS, embedded in NEG-50™ OCT compound, frozen on dry ice, and cryosectioned at 35 µm thickness. The effect of mRNA concentration on GFP transfection efficiency was assessed using one-way ANOVA followed by Sidak’s post-hoc test in GraphPad Prism.

For co-electrotransfection experiments, A5:ETE-BuSA formulated with GFP mRNA and A5:ETE-BuSA formulated with mCherry mRNA were prepared separately and mixed at a 1:1 (v/v) ratio prior to injection. For testing the effect of mRNA concentration in transfection the relative mRNA ratio was varied to 1:0.5, 1:1, and 1:1.5 (v/v). Injection, electrofunctionalization, electrotransfection, and tissue processing were performed as described above. GFP-positive, mCherry-positive, and double-positive cell populations were quantified by manual cell counting across fluorescence channels, with total cell number determined by DAPI nuclear staining. Transfection efficiency for each channel was expressed as a fraction of DAPI-positive cells across three independent experiments (n = 3), and data were visualized as grouped bar graphs with mean ± SD in GraphPad Prism.

### Gene electrotransfer in U87 Tumors

As previously described ^13,16^, fertilized chicken eggs (Gallus gallus) were incubated at 37.5°C and 65 % relative humidity in a Chicti egg incubator with automated rotation for 30 seconds, 12 times per day. The day of incubation was designated embryonic development day 0 (EDD0). On EDD4, a 2 cm-wide opening was created in the eggshell above the air sac using a mechanical egg opener to expose the chorioallantoic membrane (CAM), sealed with a 30 mm Petri dish lid, and eggs were transferred to a Heka-Brutgeräte incubator maintained at 37.5°C and 60 % relative humidity until further use. To establish an in vivo tumor model, 1.5 × 10⁶ U87 human glioblastoma cells suspended in DMEM were mixed 1:1 (v/v) with Matrigel and implanted onto the CAM on EDD9 at a site overlying large, highly branched blood vessels. Tumor growth was subsequently monitored until EDD17. On EDD 17, a Pd/Pt-coated glass capillary (30 µm diameter, BM100T- 15, Biomedical Instruments GmbH; 50 nm Pd/Pt layer, QT150 sputter coater, Quorum Technologies) loaded with 10 µL of A5:ETE-BuSA formulated with mCherry mRNA was advanced approximately 5 mm into the tumor tissue using a micromanipulator. Injection was performed at a balance pressure of 2.0 psi and an injection pressure of 8.0 psi for 5 ms (Warner PLI-100A). Following capillary retraction, an additional depot of material was deposited on the tumor surface to serve as a connection point for subsequent electrofunctionalization. Electrotransfection was performed using a Grass stimulator (Astro-Med) with the same electrode configuration. Electrotransfection was performed using a Grass stimulator (Astro-Med) with the same electrode configuration, using 60 V square pulses of 200 μs, 2 ms, 20 ms, or 200 ms duration, and 90 V square pulses of 2 ms or 20 ms duration, all at 1 Hz for 20 s (see Results for parameter optimization). Following electrotransfection, eggs were incubated for 24 hs. Tumors were then harvested and fixed overnight in 4 % paraformaldehyde in 0.1 M phosphate buffer, rinsed in PBS, cryoprotected in 25 % sucrose in PBS, embedded in NEG-50™ OCT compound, frozen on dry ice, and cryosectioned at 16 µm. Sections were mounted with ProLong™ Gold antifade reagent containing DAPI and imaged on an Olympus IX73 microscope equipped with a Hamamatsu Orca R2 camera. Flow cytometry was performed on dissociated tumor samples in parallel with immunofluorescence imaging across all electrotransfection conditions (n = 3 per condition). The harvested tumors were dissociated into single-cell suspensions by mechanical disruption through a 40 µm cell strainer using a syringe plunger. Cells were washed in FACS buffer and fixed in 2 % paraformaldehyde on ice for 15 mins, then permeabilized with 0.1 % Triton X-100 in PBS for 10 mins at room temperature, followed by two washes in FACS buffer. Cells were incubated with a rabbit anti-GFP primary antibody (1:100) for 30 mins on ice, washed, and subsequently incubated with an Alexa Fluor Plus 647-conjugated goat anti-rabbit IgG secondary antibody (1:500) for 30 mins on ice. Following two final washes, cells were resuspended in FACS buffer and immediately analyzed. Data were acquired on a BD Symphony A1 flow cytometer and analyzed using FlowJo (v10.10.0).

### In vitro ROS Detection Studies

To assess reactive oxygen species (ROS) generation under gene electrotransfer conditions, metal electrodes were compared with an in situ-formed electrode in an agarose model. In both setups, the agarose block consisted of 0.5 % agarose prepared in zebrafish Ringer’s solution supplemented with 5 µM phenol red as a colorimetric redox/pH-sensitive indicator. An Ag/AgCl electrode placed on the agarose surface served as the return electrode. An Au-coated tungsten electrode was inserted directly into the agarose block and served as the point-source electrode (anode). In the second setup, 3 µL of an A5:ETE-BuSA solution was first injected into the agarose block using a Hamilton syringe. An Au-coated W electrode connected to the resulting A5:ETE-BuSA structure then served as the working electrode (anode). Prior to pulse delivery, electrofunctionalization of ETE-BuSA onto the A5 scaffold was carried out. Square voltage pulses were subsequently delivered at 60 V or 90 V across four pulse durations (200 µs, 2 ms, 20 ms, and 200 ms) at a frequency of 1 Hz for 20 seconds.

### Electrotransfection studies with ex vivo chicken embryonic hearts

On embryonic day 17 (EDD17), chicken embryos were humanely euthanized by cervical transection. The hearts were dissected and transferred to a Petri dish containing pre-warmed, 37 °C Howard’s Ringer saline. Subsequently, the hearts were mounted on a modified Langendorff perfusion apparatus and perfused for 15 mins prior to the procedure to stabilize. Retrograde perfusion was performed through the aorta using Krebs–Ringer solution (37 °C, pH 7.4), pre- gassed with 95 % O₂ and 5 % CO₂, and delivered using a Julabo F-25 pump.The aorta was cannulated under a dissection microscope using a custom-made cannula fashioned from a 25- gauge needle. In avian embryos, the aorta branches only a few millimeters above the aortic root, requiring cannulation on the ascending aorta (Figure 1). The first two aortic branches (right and left brachiocephalic arteries) were ligated to prevent solution leakage during perfusion. Following cannulation, the heart was connected to a custom-built perfusion circuit and retrogradely perfused at a constant flow rate with Krebs solution, continuously bubbled with carbogen (95 % O₂ /5 % CO₂) at 37 °C. Hearts were maintained in a heated, humidified chamber, with temperature and hydration regulated through the perfusion cannula. Perfusion pressure was maintained at approximately 80 mmHg (range: 80–90 mmHg). Epicardial delivery was performed by depositing 15 µL of the A5:ETE-BuSA + mRNA pre-drape solution directly onto the ventricular apex using a 28-gauge Hamilton syringe. After electrofunctionalization process, electrotransfection was performed using square voltage pulses at 60 V, 1 Hz, and a 2 ms pulse width for 20 s, with a Grass stimulator (Astro-Med) and an AgCl counter electrode. Hearts were subsequently returned to the Langendorff perfusion apparatus and maintained under continuous physiological perfusion for 4 h to preserve contractile function and tissue viability. GFP-positive cell populations were assessed by flow cytometry. Treated hearts were dissected in DMEM and enzymatically digested with 450 U/mL Collagenase Type I for 20 mins at 37 °C. Remaining tissue fragments were mechanically dissociated by passage through a 40 µm cell strainer using a syringe plunger to obtain a single-cell suspension. Cells were washed in FACS buffer and fixed in 2 % paraformaldehyde on ice for 15 min, then permeabilized with 0.1 % Triton X-100 in PBS for 10 mins at room temperature, followed by two washes in FACS buffer. Cells were incubated with a rabbit anti-GFP primary antibody (1:100) for 30 mins on ice, washed, and subsequently incubated with an Alexa Fluor Plus 647-conjugated goat anti-rabbit IgG secondary antibody (1:500) for 30 mins on ice. Following two final washes, cells were resuspended in FACS buffer and immediately analyzed. Data were acquired on a BD Symphony A1 flow cytometer and analyzed using FlowJo (v10.10.0).

**Figure 1.**
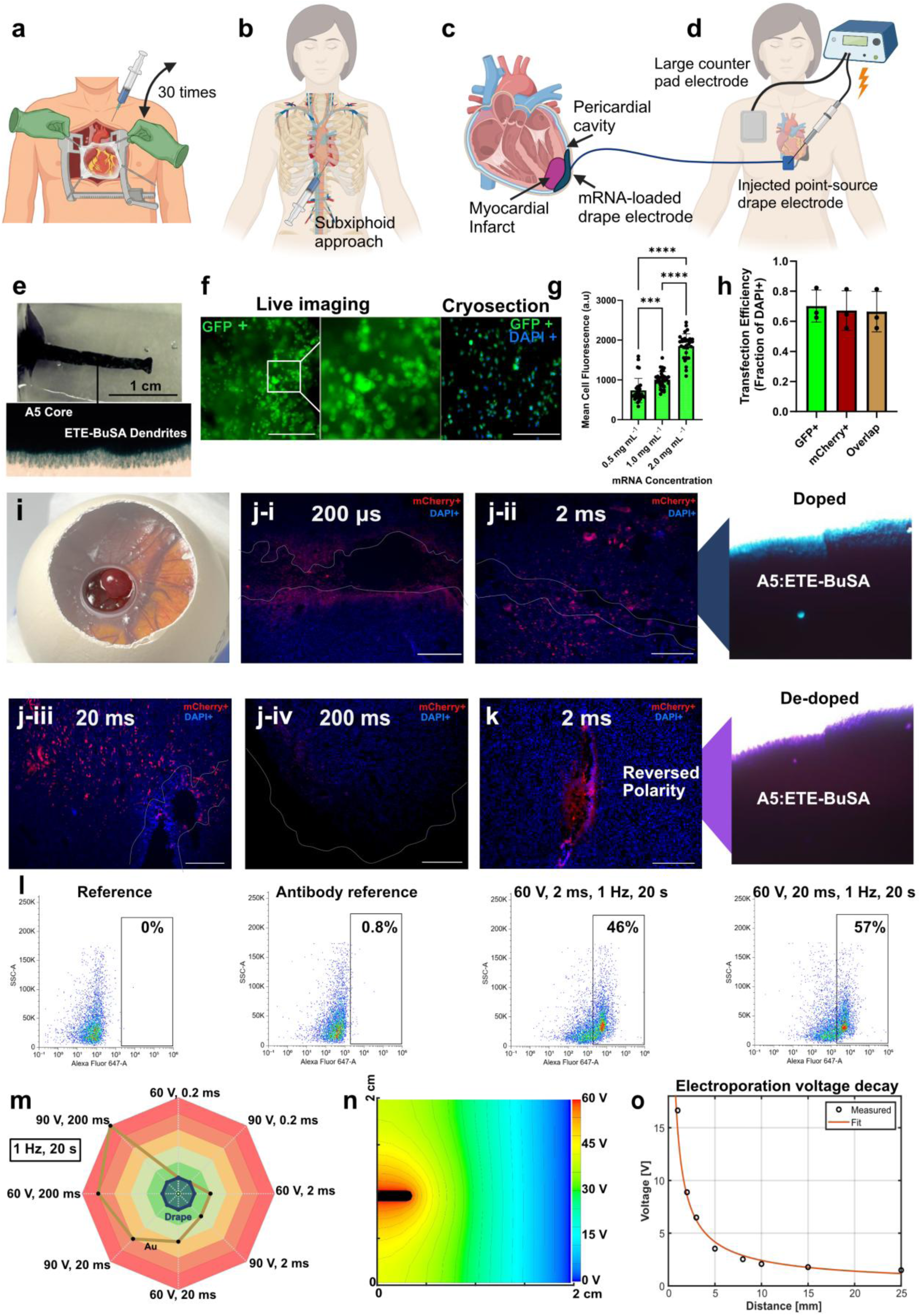
Intrapericardial mRNA delivery and electrotransfection using the A5:ETE-BuSA nanoparticle platform, validated across clinical delivery routes, 3D agarose models, and an ex ovo CAM tumor model. (a–b) Intrapericardial delivery routes: (a) open-chest surgical delivery (EPICCURE trial) vs (b) minimally invasive subxiphoid pericardiocentesis. (c) Intrapericardial injection of mRNA-loaded A5:ETE-BuSA. (d) Electrotransfection setup with point-source drape and return electrodes. (e–f) Electrotransfection of GFP mRNA–A5:ETE-BuSA in 3D agarose cell blocks: (e) Setup in agarose, (f) live fluorescence before and after cryosection. (g) Mean cell fluorescence vs. mRNA concentration (0.5–1.5 mg mL⁻¹; one-way ANOVA, Šidák’s test, ****p<0.0001, ***p<0.001, n=30–33/group). (h) Spatial co-expression overlap, GFP/mCherry. (i) U87 tumor, chicken CAM model. (j) mCherry expression in U87 CAM tumors after electrotransfection (200 µs–200 ms pulses, i–iv; 60 V, 1 Hz, 20 s; normal polarity, doped state). (k) Reversed-polarity (cathode) drape configuration (de-doped polymer). (l) Flow cytometry, GFP⁺ dissociated U87 cells: controls vs. 2 ms/20 ms pulses (60 V, 1 Hz, 20 s). (m) ROS generation (phenol red assay) of the drape electrode (deep blue enclosed area) across pulse parameters (60/90 V; 200 µs–200 ms, 1 Hz, 20 s) vs. Au electrode (gold enclosed area). (n) FEM of electric field distribution (Quickfield). (o) Peak voltage vs. distance from injection site.

### Thermal Safety Assessment by Infrared Thermography

To evaluate thermal development at the electrode–tissue interface during pulse delivery, infrared thermographic imaging was performed during electroporation of ex vivo perfused chicken hearts using a FLIR P620 infrared camera. The camera has a resolution of 640 × 480 pixels and operates in the long-wave infrared spectral range of 7.5–13 µm. It was mounted on a tripod to minimize movement artifacts, and all measurements were performed at room temperature. The apical surface of each heart was cast with the polymer drape electrode, while an Au-coated electrode contacted the drape electrode surface without direct contact with the cardiac tissue. An AgCl electrode served as the counter electrode. Temperature measurements were acquired at the interface between the drape electrode and the cardiac tissue surface before and during pulse delivery. Electrical pulses were delivered at two voltage levels (60 V and 90 V) combined with two pulse durations (2 ms and 20 ms) at a frequency of 1 Hz for 20 seconds, yielding four parameter combinations in total. Each parameter combination was tested in at least three independent experiments using three different hearts.

### Ex vivo Porcine heart gene electrotransfection

Freshly excised porcine hearts with intact pericardium were used as an ex vivo model to assess A5:ETE-BuSA-mediated intrapericardial mRNA electrotransfection. Immediately following dissection, hearts were perfused and transported in ice-cold St. Thomas’ Hospital cardioplegic solution until use. A 15 µL volume of A5:ETE-BuSA+mCherry mRNA was injected into the pericardial cavity with a Hamilton syringe, forming a localized patch at the injection site. Electrofunctionalization and electrotransfection was then performed with the same parameters used for the transfection of ex vivo chicken embryonic hearts.

### In vivo zebrafish gene electrotransfection

Adult zebrafish were anesthetized in 1.5 mM tricaine (ethyl 3-aminobenzoate methanesulfonate; Sigma-Aldrich, E10521) until cessation of opercular movements and absence of response to tail pinch, as previously described⁸. Anesthetized animals were immobilized on a sponge and mounted onto the microinjection apparatus. Depending on body size, two to four scales overlying the cardiac region were carefully removed, and a small incision was created using a 30 G Terumo Agani™ needle (0.3 × 13 mm, AN*3013R1). Injections were performed using metal-coated glass capillaries with a beveled tip of approximately 30 µm in diameter (BM100T-15; beveled, straight, shortened, and fire-polished; BioMedical Instruments GmbH), one end of which was sputter- coated in-house with a 50 nm palladium/platinum (Pd/Pt) alloy (Quorum QT150). Capillaries were loaded with 2 µL of A5:ETE-BuSA formulated with mCherry mRNA using a 20 µL Microloader tip (epT.I.P.S., Eppendorf). A micromanipulator (Quad, Sutter Instrument) was used to advance the loaded capillary through the preformed incision, oriented parallel to the sagittal plane, into the pericardial cavity. Upon confirmed entry, a single injection was delivered at a balance pressure of 1.8 psi and an injection pressure of 8.0 psi for 10 ms (Warner PLI-100A). The sustained balance pressure during capillary withdrawal resulted in deposition of an A5:ETE-BuSA layer at the skin surface, which was subsequently electrofunctionalized by applying 1.0 V through the Pd/Pt-coated capillary.

Electroporation was performed following electrofunctionalization using the same Pd/Pt-coated capillary connected to the point-source hydrogel electrode and a large AgCl electrode placed in contact with the zebrafish body as the return electrode. Square-wave pulses were delivered at 60 V and 1 Hz for 20 seconds, with pulse widths of 200 µs, 2 ms, or 20 ms, tested across experimental groups. Survival at 1 day post-injection (dpi) was assessed across the three pulse-width groups: 200 µs (n = 20), 2 ms (n = 45), and 20 ms (n = 11). Animals that exhibited overt stress responses following electroporation were humanely sacrificed. Survival proportions between the 200 µs and 2 ms groups were compared using Fisher’s exact test. Following the procedure, zebrafish hearts were collected and fixed in 2% paraformaldehyde overnight at 4°C, cryoprotected in 30% sucrose in PBS overnight at 4°C, embedded in OCT compound, and cryosectioned at 16 µm thickness. Sections were mounted onto SuperFrost Plus glass slides and stored at −20°C until further processing. To evaluate transfection efficiency, A5:ETE-BuSA formulations loaded with GFP or mCherry mRNA were tested at concentrations of 0.5, 1.0, and 1.5 µg/mL across zebrafish, chicken, and porcine cardiac models. Reporter gene expression was assessed by fluorescence microscopy on cryosectioned tissue samples.

Image analysis was performed in Fiji/ImageJ on 16-bit TIFF images. Background subtraction was applied prior to thresholding. Otsu’s thresholding was performed on a duplicate image to define intensity boundaries without altering the original 16-bit data, since applying the threshold directly converts images to binary and corrupts the mean gray value. ROIs were generated using Analyze Particles following Otsu’s thresholding on a duplicate image, with manually defined minimum and maximum particle size constraints applied to exclude background signal and non-myocardial regions. Mean gray value was recorded from the original 16-bit image in all cases. Results were normalized by subtracting the mean gray value of the control group from that of each experimental group. Statistical analyses were performed using GraphPad Prism and OriginPro. The effect of pulse duration on cardiac mCherry expression was assessed using Welch’s t-test, applied in place of the standard Student’s t-test due to unequal variances between groups. Data are presented as mean ± SD.

### Generation of MI model via left anterior descending coronary artery (LAD) ligation

Three-month-old male wild-type C57BL/6N mice were obtained from Taconic Biosciences (Ejby, Denmark) and housed in a temperature-controlled facility under a 12 h light/12 h dark cycle in groups of three to five animals per individually ventilated cage with environmental enrichment. Animals were fed standard chow and had *ad libitum* access to food and water. Upon arrival, mice were allowed to acclimate for at least 5 days before any handling or experimental procedures. Myocardial infarction was induced by permanent ligation of the left anterior descending (LAD) coronary artery as previously described^17,18^. Briefly, mice were randomly allocated and anesthetized with 2—3 % isoflurane in air. Animals were orotracheally intubated with a 22-gauge angiocatheter and mechanically ventilated at 120 breaths per minute, with a tidal volume of 200- –250 µL and a positive end-expiratory pressure of 3 cm H₂O (MiniVent, Hugo Sachs, March, Germany). A left lateral thoracotomy was performed to expose the heart, after which the pericardium was opened, and the left anterior descending (LAD) artery was permanently ligated using a 7-0 non-absorbable suture. The chest was subsequently closed in layers, and animals were extubated following return of spontaneous breathing. Mice received pain medication (2 µl/g mouse buprenorphine 0.05 mg/ml via intraperitoneal injection) for up to three days post-surgery. This model shows stable cardiac injury 6 weeks after MI^19^. Sample sizes were determined based on prior studies using this model^18,20,21^, and subsequent analyses were performed by investigators blinded to group allocation.

### Echocardiography aided in vivo transthoracic injections of A5:ETE-BuSA drape electrode

Mice with successful MI underwent echocardiographic assessment and echo-guided left ventricular (LV) injection at predefined time points after infarction. Mice were anesthetized with 1.2 % isoflurane (Forene, AbbVie) delivered by continuous inhalation with artificial ventilation throughout the procedure. Chest hair was removed using depilatory gel, and animals were placed on a heating pad with their limbs connected to ECG electrodes. Echocardiography was performed using a Vevo 2100 imaging system equipped with an ultra-high-frequency linear array transducer (MS550D; VisualSonics). Parasternal long-axis cine loops were acquired at >1,000 frames s⁻¹ using ECG-gated kilohertz visualization, and parasternal short-axis cine loops were obtained at 1, 3, and 5 mm below the mitral annulus. All experiments were completed within 30 mins after anesthesia induction. For echo-guided LV injection, an injection site above the infarct zone was selected from parasternal short-axis images. Under real-time B-mode imaging guidance, the needle was advanced into the LV cavity. A 30 G needle attached to a Hamilton injector was used for all injections. Each animal received 7 µL of A5:ETE-BuSA. Care was taken to avoid air bubbles during loading and injection. Following delivery, electrofunctionalization, and electroporation, the needle was withdrawn, and gentle pressure was briefly applied at the insertion site to minimize bleeding. Animals were then allowed to recover on a heating pad and monitored until spontaneous movement resumed. Data were analyzed offline in a blinded manner using VevoLab software (VisualSonics).

### In vivo intervention with A5:ETE-BuSA drape point-source electrode

One week after LAD ligation, a micromanipulator-mounted 28 G Hamilton syringe was used to inject the VEGF-A mRNA-loaded A5:ETE-BuSA drape electrode into the infarct zone via left lateral thoracotomy, exposing the heart with the pericardium left intact. The formulation was electrofunctionalized at 1 V for 3 mins, and gene electrotransfer was performed in vivo using square-wave pulses (2 ms pulse width, 1 Hz, 20 s). To restrict electrical contact to the injection tip, the Hamilton needle was coated with Pd/Pt at the tip, while the remaining surface was insulated with nail polish. Mice were randomly assigned randomized into one of two groups: vehicle (drape electrode without mRNA: A5:ETE-BuSA), and drape electrode with VEGF-A mRNA (A5:ETE- BuSA+ VEGF-A mRNA) formulation. The surgeon performing the procedures was blinded to group allocation. Post-operative analgesia was provided with buprenorphine (0.05 mg kg^-1^, subcutaneously every 12 h for 2–4 doses).

Cardiac function following each intervention was evaluated by cardiac MRI using a 9.4 T preclinical scanner combining an Agilent magnet with Bruker BioSpec Avance III electronics, operated under ParaVision 7.0.0 as previously described^20^. The system was equipped with actively shielded gradients incorporating integrated shims (BGA 12S HP, Bruker; maximum gradient strength 670 mT m^-1^; rise time 130 µs). Cardiac imaging was performed using a modified Bruker 2D ultra-short echo time (UTE) sequence with the following parameters: echo time 0.5 ms, repetition time 5 ms, slice thickness 1 mm, matrix size 96 × 96, field of view 25 × 25 mm², flip angle 7°, and 12,064 spokes per slice acquired within 1 minute. Image-based determination of EF, stroke volume, cardiac output, end-diastolic volume, end-systolic volume, and left ventricular mass was performed with Segment (Medviso, Lund, Sweden)^22^. Regional wall assessment was conducted on a slice-by-slice basis, following the previously described^23^methodology. Briefly, the LV was represented by five short-axis slices, ranging from the apical region to the basal region. Each slice was further subdivided into six sectors. Based on the qualitative evaluation of the bulls- eye plots, myocardial scar tissue was predominantly observed in slices 3–5. Therefore, these three slices were selected for subsequent quantitative regional wall analysis. Quantitative multiplanar measurements were then performed for the corresponding septal and lateral sectors.

For MRI contrast-enhanced visualization of the formed drape electrode, A5:ETE-BuSA was prepared for MRI contrast-enhanced visualization by dissolving ETE-BuSA at 40 mg mL^-1^ in 1 mM Gadoteridol solution (Cayman Chemicals, Item #: 33577) in deionized water, followed by dissolution of A5 at 20 mg mL^-1^ in the same solution to yield the final formulation. In a separate cohort, one week after LAD ligation, the Gadoteridol-carrying A5:ETE-BuSA was delivered into the infarct zone via a micromanipulator-mounted 28 G Hamilton syringe under left lateral thoracotomy. Electrofunctionalization and electroporation were subsequently performed as described above.

### Protein isolation

Heart tissue sectioned to left ventricle (LV), right ventricle (RV) and septum and LV samples were homogenized in RIPA lysis buffer (150 mM NaCl, 50 mM Tris-HCl, pH 8.0, 2 mM EDTA, 2% Triton X-100, 0.05 % SDS, and 0.5 % sodium deoxycholate) supplemented with protease and phosphatase inhibitor cocktails. Lysates were incubated on ice for 30 mins with intermittent vortexing and then sonicated (10 % amplitude, 0.5 s pulse, 10 s total). Samples were centrifuged at 16,000 × g for 15 mins at 4°C, and the supernatants were collected. Protein concentrations were determined using the bicinchoninic acid (BCA) assay. Equal amounts of protein (9 mg mL^-1^) were mixed with 1× RIPA, and subsequently 25 µL of the mix was used for collagen quantification.

### Collagen quantification in left ventricle homogenates

Total collagen content in cardiac tissue was determined using a Sirius Red-based colorimetric assay. RIPA extracts from LV tissue 0.25 µL sample was transferred into a 96-well plate in duplicate and kept in an incubator at 37°C overnight to allow it to dry. The samples were fixed in 150 µL of Bouin’s solution (consisting of 75 % saturated picric acid, 25 % formaldehyde, and 5 % glacial acetic acid) for one h at room temperature. The wells were then washed three times with distilled water and incubated for 1 h at room temperature with 150 µL of Sirius Red solution (1 g L^-1^ Sirius Red F3B (Alfa Aesar, B21693) in saturated picric acid). After staining, the wells were washed three times with distilled water to remove the unbound dye. The bound Sirius Red was then eluted with 150 µL of 0.1 M NaOH for 30 mins at room temperature with gentle shaking (200 rpm). Then, 120 µL of the extracted dye solution was transferred to a new 96-well plate, and the absorbance was measured at 540 nm using a microplate reader. A collagen I standard prepared by diluting rat tail Collagen I solution (Gibco, A10483-01) with 0.1 N HCl 1:1 for the following concentrations: 3, 1.5, 1, 0.75, 0.5, 0.25, 0.125 was used as quantification control on each plate and underwent the same treatment as the samples. Collagen concentrations were calculated from the standard curve and expressed as milligrams of collagen per milliliter of tissue lysate.

### Mouse NT-proBNP ELISA

Levels of mouse NT-proBNP in cardiac tissue lysates were determined using a commercially available ELISA kit (Novus Biologicals, Centennial, CO, USA; catalog no. NBP2-76775) according to the manufacturer’s instructions. Tissue homogenates were prepared in PBS, centrifuged and the resulting supernatant was collected for analysis. Samples and standards were assayed in duplicate. Following sequential incubation with a biotinylated detection antibody and an HRP-conjugated avidin, the plates were developed using a substrate solution and the reaction was stopped using a stop solution. Absorbance was measured at 450 nm using a microplate reader. NT-proBNP concentrations were calculated from a four-parameter logistic (4-PL) standard curve and expressed in pg mL^-1^.

### Histology of mouse tissues

Following transcardiac perfusion with saline, tissues were harvested and post-fixed in 4% paraformaldehyde for 24 hs at 4°C. Specimens were then either cryoprotected in 30% sucrose in PBS (137 mmol L^-1^ NaCl, 2.7 mmol L^-1^ KCl, 10 mmol L^-1^ Na₂HPO₄, 1.8 mmol L^-1^ KH₂PO₄) and kept at 4°C, or kept at room temperature until sectioning. 10 μm cryostat-sectioned aorta cross sections were fitted onto SuperFrost Plus glass slides, washed with PBS and then sequentially incubated in Mayer’s hematoxylin (Histolab) for 10 mins, differentiated in HCl-ethanol (0.3 % in 70 % ethanol) for 1 minute, rinsed in tap water for 5 mins, and counterstained with eosin (0.5 g in 70 % ethanol) for 1 minute. Sections were dehydrated, coverslipped with Rotimount ((Carl Roth, HP68.2), and imaged on Nikon Ti2 WideField FM light microscope. For Masson trichrome staining, a commercial kit was used with minor modifications to the protocol. Briefly, sections were fixed in Bouin’s solution for 5 mins at 57 °C, rinsed under running tap water for 3 mins, and washed in distilled water. Sections were then stained with Weigert’s iron hematoxylin for 1 minute, washed under tap water for 5 mins, rinsed, and incubated in Biebrich scarlet-acid fuchsin for 45 seconds. After a 3-minute treatment with phosphotungstic/phosphomolybdic acid, sections were placed in aniline blue for 45 seconds and then in 1 % acetic acid for 1 minute. Slides were cover- slipped with Rotimount and imaged on an Olympus BX60 or CX33 microscope. 10 μm vertically heart cross sections were fitted onto SuperFrost Plus glass slides, washed with PBS and incubated for 30 mins at room temperature with blocking buffer (#11096176001, Roche, Switzerland), followed by overnight incubation with primary antibodies against anti-Vimentin (Vim; ab8978, Abcam), and FITC-Lectin (Lectin from Lycopersicon esculentum (tomato); L0401, Sigma- Aldrich) After washing in PBS, samples were incubated with Alexa Fluor conjugated secondary antibody (Goat Anti-Mouse Alexa Fluor 594 #A-11032, Invitrogen) for 1 h, washed, mounted with Fluoromount-G Mounting Medium with DAPI (#00–4959-52, Invitrogen, USA), and examined under a Nikon TiE A1RHD Confocal FM or a Leica STELLARIS 5 microscope.

### RNA isolation from LV tissue and quantitative RT-PCR

RNA was isolated using TRIzol Reagent (Thermo Fisher Scientific, Cat#15596018) according to the manufacturer’s instructions. 200 µg of mRNA was reverse transcribed into cDNA using the High-Capacity cDNA Reverse Transcription Kit (Thermo Fisher, Cat#4368814) in a T100 Thermal Cycler (Bio-Rad, Hercules, CA, USA). The resulting cDNA was diluted 1:12.5 and used as a template for quantitative RT-PCR. Gene expression (see Table S1 for primer pairs used in qPCR) was detected using SYBR Green PCR Master Mix (Thermo Fisher Scientific, #4369702) and CFX384TM Real-Time System (Biorad, Hercules, CA, USA). All samples were run in duplicates, and the gene expression relative to the housekeeping gene L14 was calculated from the standard curve using the delta Ct method.

### Cardiac data analysis

Sample sizes for murine experiments were based on prior experience with the models^17,18,24^. No animals or data points were excluded. Data from mouse experiments were analyzed using GraphPad Prism (RRID:SCR_002798; version 10). The sample size (N) denotes the number of animals; exact N values are provided in the figure legends. Data are presented as mean ± SD after evaluation of data distribution using the Shapiro-Wilk test. Comparisons between two dependent groups were performed using paired Student’s t-test. For comparisons involving multiple groups, two-way analysis of variance (ANOVA) or Repeated Measure ANOVA with LSD post-hoc testing were applied, with experimental group and time post-MI included as factors. P values < 0.05 were considered statistically significant.

### Animal Ethics

This study was conducted in accordance with Swedish national legislation and European Community guidelines for animal studies. The ethics committee in Malmö-Lund approved all procedures (5.8.18- 05993/2018, 5.8.18–08160/2021, 5.8.18-05748/2022, 5.8.18-19103/2023, 5.8.18-14704/2023 and 5.8.18-10213/2026).

## Results & Discussion

The open-chest surgical approach used in the EPICCURE study, compared with the subxiphoid injection used for pericardiocentesis, is illustrated in Figure 1a–d. Our nanoparticle solution is compatible with these clinically established intrapericardial delivery routes. The nanoparticle solution can be delivered via needles with outer diameters as small as 30 μm^25^, substantially smaller than those typically used clinically for pericardiocentesis (1.3–1.7 mm).

Our in situ-formed conductive epicardial drape is electrically distinct from conventional gene electroporation systems. Classical gene electrotransfer typically relies on nucleic acid injection administered separately, followed by pulse delivery between two or more closely spaced electrodes, with the treated volume primarily defined by the electric field established between them. In contrast, our approach uses point-source electroporation, also referred to as a monopolar configuration, in which a small epicardial active patch is paired with a distant large return electrode^12,14^. This creates a monopolar-like geometry with a strongly non-uniform electric field confined at the patch.

Importantly, in our case, the point-source electrode serves a dual function: it acts both for the local mRNA delivery and as the active electrode for pulse delivery. Because mRNA is released from the same patch that generates the highest local electric field, nucleic acid availability spatially coincides with the region of most efficient electroporation, favoring localized epicardial myocardial transfection.

A key initial question was whether incorporation of mRNA into the polymer mixture would preserve the self-assembly and electrochemical properties of the A5:ETE-BuSA dispersion used for BICS. GFP reporter mRNA was mixed into the polymer dispersion and injected into a U87 cell-laden agarose model, in which U87 cells serve as a cellular target and the agarose matrix recreates a three-dimensional, tissue-like environment. Following injection of the A5:ETE-BuSA+ mRNA formulation into the agarose block, the polymer mixture underwent non-externally triggered self-assembly, and the subsequent electrofunctionalization generated dendritic structures by volumetrically incorporating ETE-BuSA trimers onto the A5 conductive scaffold, consistent with our previous observations (Figure 1e). To further assess whether the mRNA cargo affected the formed hydrogel electrical properties, electrochemical impedance spectroscopy (EIS) was performed with and without mRNA. The impedance spectra largely overlapped over the measured frequency range (Figure S1), indicating that mRNA incorporation has little to no effect on the electrode’s electrochemical behavior or conductivity.

Gene electrotransfection enabled efficient delivery of the mRNA payload into U87 cells, as demonstrated by GFP expression (Figure 1f). Quantification of mean fluorescence intensity (MFI) across ROIs from three biological replicates showed a dose-dependent increase in transfection: MFI values increased with mRNA concentration, reaching 322.1, 638.9, and 1105 at 0.5, 1.0, and 1.5 mg mL⁻¹, respectively (Figure 1g). All pairwise comparisons were statistically significant, indicating that transfection efficiency scales close to linearly with mRNA concentration within this range. Importantly, cells treated with only A5:ETE-BuSA showed negligible background fluorescence, confirming that the observed expression was attributable to mRNA delivery rather than autofluorescence or nonspecific signal.

To assess the delivery versatility using the electroactive device, GFP- and mCherry-encoding mRNAs were mixed in equal ratios in the A5:ETE-BuSA formulation and co-delivered into U87 cell-laden agarose blocks (Figure 1h, Figure S2). Fluorescence imaging showed robust expression of both reporters, with approximately 70% spatial overlap between the GFP and mCherry signals, indicating that a substantial fraction of transfected cells received and expressed both mRNA species simultaneously. This level of co-expression is notable because it was achieved from a single injection point within a 3D matrix, without additional targeting strategies or sequential delivery steps, indicating the flexibility of the electroactive device for multiplexed mRNA delivery.

To further optimize electroporation parameters, A5:ETE-BuSA+mCherry mRNA was injected into U87 tumors on the chicken CAM model (Figure 1I) and electrotransfected across a range of pulse durations at 60 V (200 µs to 200 ms; Figure 1j i–iv). These parameters were selected based on previous studies in swine and mice. Pulse duration strongly influenced both transfection efficiency and the distribution of reporter expression in the tumor. With 200 µs pulses (Figure 1J- i), mCherry expression was barely detectable and restricted to the immediate vicinity of the injection site, consistent with limited effective field penetration into the surrounding tumor tissue. Increasing the pulse duration to 2 ms (Figure 1j-ii) resulted in a broader distribution and more homogeneous expression across the tumor slices, indicating efficient electroporation of cells located farther from the injection point. A comparable expression pattern was maintained at 20 ms (Figure 1j-iii).

In contrast, at 200 ms (Figure 1j-iv), reporter expression was largely absent, consistent with a shift toward irreversible electroporation and loss of cell viability before transgene expression could occur. Thus, pulse duration defines a critical operational window: pulses that are too short limit tissue-wide gene delivery, whereas excessively long pulses become cytotoxic.

The colorimetric transition from blue to purple reflects changes in the redox state of the conducting polymer. In the doped, oxidized state, charge carriers are introduced along the polymer backbone, giving the material its blue appearance. Upon reduction and de-doping, these charge carriers are depleted, shifting the transmitted color toward purple, as observed in the reversed-polarity condition, where cathodic bias de-doped the polymer and minor mCherry expression was detected only very close to the electrode.

To optimize the electroporation parameters for transfection efficiency while avoiding irreversible tissue damage, we tested combinations of voltage and pulse duration beyond the 60 V, 2 ms setting. The imaging data (Figure 1j) were complemented by quantitative flow cytometric analysis of, dissociated U87 tumors (Figure 1l). At 60 V, a 2 ms pulse resulted in 46.1% GFP-positive cells, while extending the pulse duration to 20 ms produced a comparable positive fraction of 55.9%. Increasing the voltage to 90 V at 2 ms yielded a similar transfection efficiency (54.0%) (Figure S3), indicating that voltage alone within this range did not substantially alter the transfection outcome. In contrast, combining the higher voltage with the longer pulse duration (90 V, 20 ms) reduced the GFP-positive fraction to 23.4% (Figure S3).

PEDOT:PSS-coated metal electrodes have previously been shown to improve cell viability and transfection efficiency compared with bare metal electrodes, such as Au, likely because the volumetric capacitance of the polymer reduces interfacial electron transfer and thereby limits reactive oxygen species (ROS) generation in the tissue. We therefore evaluated whether our free- standing, substrate-free electrode generated ROS across the tested parameters, compared with an Au-coated electrode, using phenol red as a colorimetric readout in agarose (Figure 1m). No ROS reactions were detected using the tested parameters with our drape electrode (dark blue within the green area, Figure 1m). However, with an Au-coated electrode, ROS development was observed at the longer pulse lengths of 20 ms and 200 ms at both 60 and 90 V. These findings are consistent with previous studies^26^. Notably, even when the voltage was increased well beyond 90 V, up to 175 V, under both anodic and cathodic stimulation conditions, no ROS were detected.

Point-source electroporation in a monopolar configuration has been described in previous modeling studies, and our experimental data corroborated the predicted field distribution and treatment behavior. Thus, finite element modeling of the point-source geometry in Quickfield. Multiphysics showed a steep voltage gradient radiating from the injection site, with field intensity falling sharply over distance (Figure 1n). The model showed the field concentrated at the electrode, with the steep-gradient, high-field region relevant to electroporation confined to within roughly 2– 3 mm of the injection site. Although the potential extends further into the surrounding tissue, the gradient there is shallow, so the field falls below the permeabilization range, and the effective electroporation zone remains localized near the electrode. Experimental, direct peak-voltage measurements reproduced this profile (Figure 1o), decaying from roughly 17 V at the site to under 2 V at 15 mm and to negligible values beyond 25 mm. The measured potential can be fitted using a power law with an exponent of −0.788. This is close to the −1 expected for an ideal point-source electrode in a homogeneous conductive medium^27^. The electric field, and therefore the electroporation effect, is confined to the immediate surroundings of the injectable electrode, leaving distant tissue largely unstimulated. Together, pulse duration and electrode geometry constrain delivery along two complementary axes: pulse duration defines the intensity of electroporation, while electrode geometry defines its spatial reach. This provides control over both how strongly and how far the field acts within the tissue.

Our injectable liquid-electrode technology is therefore particularly well suited for point-source electroporation. Unlike rigid electrodes, the material can conform to the local tissue architecture or, as previously described, be delivered by convection-enhanced delivery to occupy a defined tissue volume. Because electrode geometry determines the electric-field distribution, the liquid electrode directly governs the spatial extent and pattern of electroporation.

### Ex vivo cardiac studies

Up to this point, the platform had been characterized under static conditions. The heart it is ultimately meant to treat is neither static nor quiescent: it contracts continuously and is perfused throughout, where the surface in motion and the flow across it both bear on whether the hydrogel can form, adhere, and transfect. Whether these conditions affect the platform’s formation and function is the question an ex vivo beating heart can address directly. We established Langendorff- perfused chicken embryo hearts on embryonic day 17 (EDD 17) for this purpose. At this stage, the heart is fully four-chambered with complete septation, so its architecture and the separation of its circulation resemble the mammalian and human heart far more closely than simpler two- chambered models, while remaining large and accessible enough for intrapericardial casting, electrode placement, and aortic cannulation. Retrograde perfusion through the aorta maintains coronary circulation, contraction, and metabolic activity in isolation from systemic influences, providing a controlled setting in which the local effects of an intrapericardial intervention can be resolved precisely before in vivo work. Once formed on the myocardium, the drape electrode remained adherent to the cardiac surface over 4 to 6 hs of perfusion, without visible disruption (Figure 2a).

**Figure 2.**
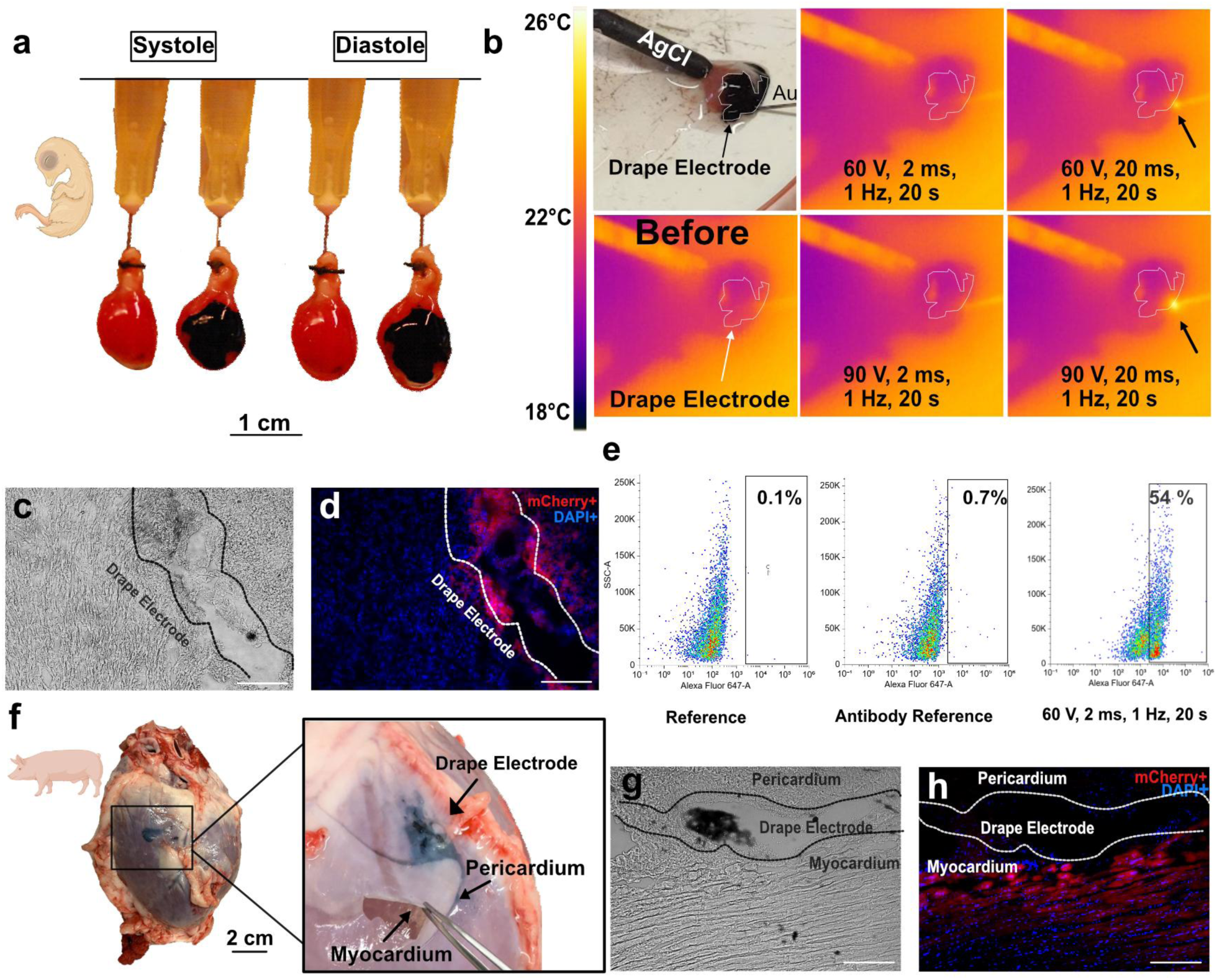
Ex vivo cardiac validation of the A5:ETE-BuSA mRNA delivery platform in chicken embryo and porcine heart models. (a) Langendorff-perfused chicken embryo hearts (EDD17) presented with A5:ETE-BuSA drape electrode at apical surface after epicardial deposition/electrofunctionalization within systolic and diastolic phases. (b) IR thermography at the drape electrode–tissue interface, before vs. during pulsing, across four electrotransfection parameter combinations. (c–d) Apical myocardium cryosection after intrapericardial electrotransfection (mCherry mRNA–A5:ETE-BuSA): (c) brightfield, (d) corresponding mCherry fluorescence. (e) Flow cytometry, GFP⁺ fraction in dissociated Langendorff heart cells (60 V, 2 ms, 1 Hz, 20 s). (f) Ex vivo porcine heart after intrapericardial A5:ETE-BuSA injection: hydrogel confined within pericardial cavity; opened pericardium showing gel-like drape electrode. (g) Histology, porcine cardiac tissue: hydrogel within pericardial space at injection site. (h) Immunofluorescence, porcine cardiac cryosection after electrotransfection (mCherry mRNA– A5:ETE-BuSA): mCherry⁺ perimyocardial tissue.

Thermal injury is a critical safety consideration in electroporation-based cardiac interventions, and FDA guidance on thermal effects of medical devices explicitly includes electroporation-based systems within its evaluation framework. In cardiac tissue, temperatures of 50–55°C are generally associated with irreversible myocardial injury, although the precise threshold depends on the duration of exposure and the cumulative thermal dose^28^. Sufficiently high voltage or prolonged pulses can in principle drive localized Joule heating at the electrode-tissue interface, which would threaten both tissue integrity and mRNA stability. To test for this, the drape electrode-tissue interface was imaged by infrared thermography during electrotransfection of ex vivo perfused chicken hearts across four parameter combinations (Figure 2b). At 60 V / 2 ms no temperature increase was detected (ΔT = 0 °C), and at 90 V / 2 ms the rise was negligible (0.3 °C). Extending the pulse to 20 ms produced larger but still modest effects, with ΔT of 1.6 °C at 60 V and 3.2 °C at 90 V, accompanied by localized heating at the interface. The maximum increase of 3.2 °C remained well below the damage threshold, placing all tested conditions within a thermally safe window. This reinforces 2 ms as the working pulse duration, combining effective delivery with a negligible thermal increase.

With the drape electrode conforming to the cardiac surface on a beating, perfused heart and the protocol confirmed to remain within a thermally safe window, the remaining question was whether transfection would proceed under these conditions, where continuous motion and coronary flow could limit it. Cryosections of the apical myocardium showed detectable mCherry fluorescence within the tissue (Figure 2c and 2d), indicating that delivery and expression occur despite the mechanical activity and coronary flow of the perfused heart. Thus, the drape electrode adheres to the beating heart, delivers mRNA, and generates an electric field that enables electrotransfection of the underlying myocardium during physiological motion.

Quantifying the expression by flow cytometry on the dissociated chicken embryo hearts, the optimized parameters (60 V, 2 ms, 1 Hz, 20 s) yielded GFP-positive cells in approximately 54% of the population, against background fractions of 0.1% and 0.7% in the reference and antibody- reference gating controls, respectively (Figure 2e).

We next turned to the porcine heart, which resembles the human heart anatomically. Following intrapericardial injection, the assembled drape electrode was visible within the pericardial space as a coherent structure on the pericardium lining the epicardial surface (Figure 2f). Upon exposure of the pericardium, the A5:ETE-BuSA+mRNA drape electrode exhibited a gel-like consistency, similar to pericardial fluid, and could move in concert with the heart. This mechanical behavior is consistent with the tissue-compatible elastic Young’s modulus of 21 kPa previously reported for eBICS^13^, confirming that the formulation retains its structural and mechanical integrity within the pericardial space of a large animal heart without disrupting pericardial compliance.

Immunofluorescence examination of porcine cardiac tissue sections confirmed the presence of the deposited hydrogel within the pericardial space at the expected injection site (Figure 2g), consistent with the delivery pattern observed in smaller animal models and supporting the premise that the formulation does not migrate beyond the target site. mCherry fluorescence was detectable within the pericardial and myocardial tissue following electrotransfection (Figure 2h), confirming that mRNA delivery and expression occur in the porcine cardiac setting. Fluorescence intensity peaked within the 200–300 µm range of the in situ-formed drape electrode, then declined progressively beyond 300 µm and approached baseline levels beyond 500 µm (Figure S4).

### In vivo intrapericardial delivery in zebrafish

Ex vivo models, while informative, are limited in the duration over which experiments can be sustained in vivo. We turned to the zebrafish, previously used in our BICS study^13^, whose accessible pericardial space makes it uniquely suited as the first test model for intrapericardial injections under direct stereomicroscopic visualization. Intrapericardial electrode installation of the A5:ETE-BuSA + mCherry mRNA formulation was performed using a micromanipulator- assisted capillary (Figure 3 a-i), after which the formulation self-assembled within the pericardial cavity and electrofunctionalization was carried out to form the drape electrode in situ and the subsequent gene electrotransfer. Following the procedure, zebrafish were revived and monitored, exhibiting swimming and feeding patterns indistinguishable from those of untreated controls, with no signs of distress (Figure 3a-ii). Excision of the heart at endpoint confirmed that the drape electrode had assembled on the cardiac surface (Figure 3b). Thus, the mRNA cargo does not impair the drape formation in vivo. To assess in vivo tolerability, survival was monitored as a function of pulse duration following intrapericardial electrotransfection (Figure 3c). Survival rates of approximately 80% and 85% were observed at 200 µs and 2 ms, respectively, with no statistically significant difference between these conditions. Extending the pulse duration to 20 ms reduced survival to approximately 25%, indicating that 20 ms is substantially lethal.

**Figure 3.**
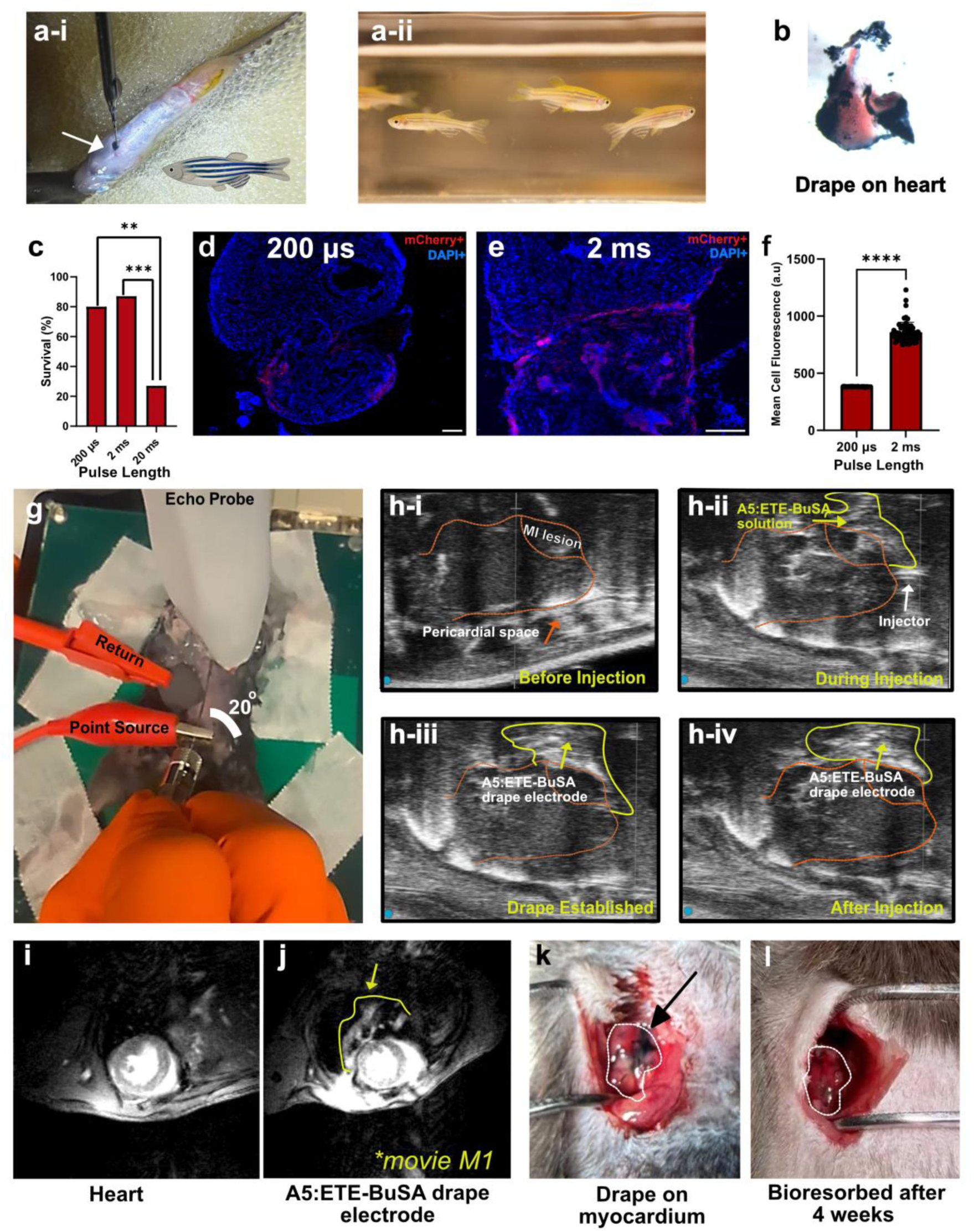
In vivo validation of intrapericardial A5:ETE-BuSA delivery and electrotransfection in zebrafish and a post-myocardial infarction mouse model. (a-i) A5:ETE-BuSA + mRNA intrapericardial injection in anesthetized zebrafish, stereomicroscopic guidance. (a-ii) Recovered zebrafish post-procedure. (b) Excised zebrafish heart, drape electrode conforming to cardiac surface. (c) Zebrafish survival across three pulse durations (n=51 fish, Fisher’s exact test). (d–e) Immunofluorescence, excised zebrafish hearts after electrotransfection: (d) 200 µs, (e) 2 ms. (f) MFI quantification, zebrafish hearts, 200 µs vs. 2 ms. (g) Echo-guided closed-chest intrapericardial injection (A5:ETE-BuSA + mRNA) in post-MI mouse (Vevo echocardiography). (h) Real-time echo sequence of delivery workflow: (i) pre-injection heart position, (ii) needle advancement/deposition, (iii) assembled drape electrode post-electrofunctionalization, (iv) stable electrode positioning post-injection. (i–j) Cardiac MRI: (i) control heart, (j) Gadoteridol-loaded drape electrode (see Supplementary Movie M1). (k) Drape electrode within pericardial space overlying MI region, post-injection/electrofunctionalization. (l) Pericardial space, four weeks post- drape electrode formation.

### Echo-guided intrapericardial delivery in mice

Although intrapericardial delivery is clinically established, we assessed its feasibility in a disease- relevant mouse model as a stringent preclinical test of the approach. This represents a greater technical challenge than in humans, owing to the smaller size of the animal and the considerably higher murine heart rate, which typically ranges from 400–600 beats per minute compared with 60–100 beats per minute in humans. The injection procedure was performed in mice one month after MI induction via LAD coronary artery ligation, a well-established model of heart failure with reduced ejection fraction^20^, allowing us to assess whether MI-associated structural changes to the pericardium and myocardium would preclude successful hydrogel formation. Injection was performed via the ultrasound-guided percutaneous subxiphoid technique^29^, in which a needle is introduced under continuous high-resolution echocardiographic guidance through the subxiphoid region into the pericardial sac without thoracotomy (Figure 3g). Injection was performed using a Pd/Pt-coated Hamilton syringe under real-time echocardiographic guidance, enabling continuous visualization of needle placement and formulation deposition throughout the procedure.

Prior to injection, the pericardial space was clearly delineated in the echocardiographic field of view (Figure 3h-i). During injection, the advancing needle and the entering A5:ETE-BuSA solution were visualized in real time, confirming accurate intrapericardial placement without inadvertent myocardial or ventricular penetration (Figure 3h-ii). Following injection, the assembled hydrogel was distinguishable as a drape electrode conforming to the cardiac surface during electrofunctionalization (Figure 3h-iii). Post-procedural imaging confirmed stable electrode positioning within the pericardial space, with the MI lesion remaining clearly identifiable throughout (Figure 3h-iv). In a preliminary cohort of three animals, successful intrapericardial deposition was confirmed in all cases, demonstrating that closed-chest echo-guided delivery remains feasible in the structurally remodeled post-infarction heart.

### Visualization of intrapericardially established drape electrode by cardiac MRI

Echocardiography guided the injection in real time and confirmed intrapericardial placement. We next examined how the assembled drape electrode behaved within the intact pericardial space during cardiac motion. Cardiac MRI visualized an assembled gadoteridol-loaded drape electrode on the heart in vivo. No signal was observed in control animals (Figure 3i), whereas the deposited material was clearly delineated within the pericardial compartment in treated animals (Figure 3j). Throughout the cardiac cycle, the material moved synchronously with the beating heart (Video M1). These MRI findings were confirmed by direct open-chest inspection, which also enabled tracking of the electrode from formation through clearance. Immediately after injection, electrofunctionalization and electroporation were performed, and the drape electrode covered the cardiac surface, including the infarct region (Figure 3k). One month later, it had largely bioresorbed, leaving only trace remnants on the cardiac surface (Figure 3l). To assess the systemic toxicity and tolerability of the material, a contract research organization (CRO) performed comprehensive hematological (16 parameters) and clinical chemistry (17 parameters) analyses, together with histopathological evaluation, in mice implanted with the electrodes. Hematology and clinical chemistry were comparable across groups and time points, and general health and body weight were unaffected.

### *Delivering VEGF-A mRNA via a drape electrode improves cardiac function after myocardial* infarction

Cardiac MRI was performed to assess the therapeutic effects of delivering VEGF-A mRNA with an A5:ETE-BuSA drape electrode system on cardiac function following MI. MRI was performed one week post-MI, before treatment initiation, and subsequently at three and six weeks post-MI, corresponding to two and five weeks after treatment, respectively. Representative cardiac MRI images at each time point are shown for the A5:ETE-BuSA and A5:ETE-BuSA + VEGF-A mRNA groups (Figure 4a&b and Supplementary Movie M2 and & M3 for 6 weeks post MI). One week after MI, before treatment, ejection fraction was reduced to 46.0 +/- 7.1 % in all MI groups (Figure 4c), as previously described^30^. Following a single administration, ejection fraction remained reduced in animals receiving the A5:ETE-BuSA drape electrode without VEGF-A mRNA, with values comparable to those of untreated MI animals (Figure S5a). In contrast, ejection fraction progressively improved in animals treated with the VEGF-A mRNA-loaded A5:ETE-BuSA drape electrode, approaching levels observed in healthy pre-MI mice by 6 weeks post-MI (Figure 4c & Figure S5a). A similar pattern was observed for stroke volume, which increased progressively following VEGF-A mRNA treatment, reaching levels comparable to the healthy pre-MI mouse group by 6 weeks post-MI (Figure 4d & Figure S5b). Importantly, administration of the A5:ETE- BuSA drape electrode alone did not produce a comparable improvement (Figure 4d), indicating that the observed functional benefit was associated with VEGF-A mRNA delivery rather than the electrode material itself^31^. Similar findings were observed for end-systolic volume (Figure 4e & Figure S5c) and cardiac output (Figure 4f), albeit without significant difference between the groups at either post-treatment time point. The findings are particularly notable given the low dose of VEGF-A mRNA delivered in a single administration, highlighting the potential of localized delivery and electrofunctionalization-assisted drape formation to achieve therapeutic effects while limiting the amount of mRNA required.

**Figure 4.**
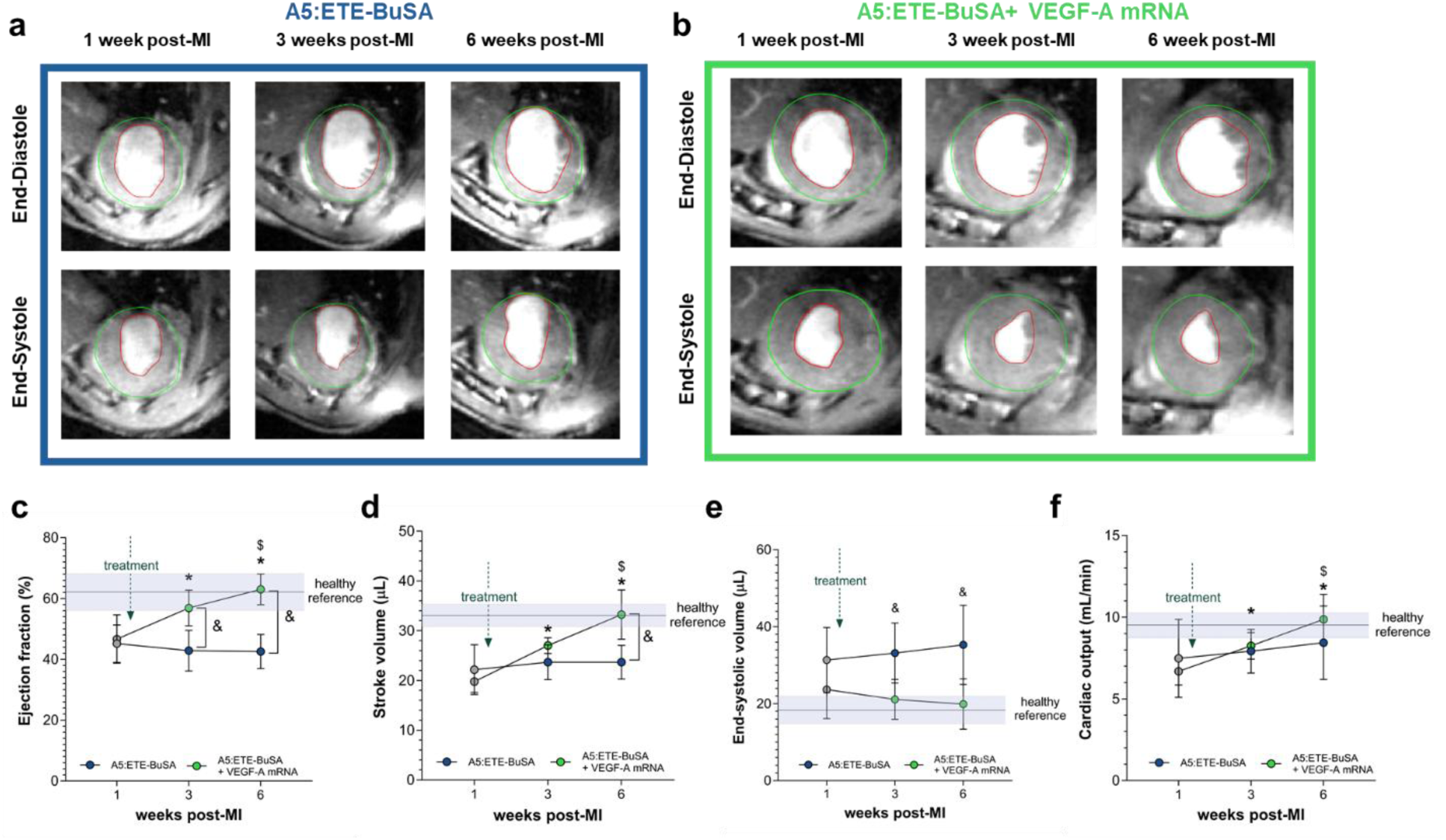
Longitudinal cardiac MRI assessment of left ventricular function following VEGF-A mRNA delivery. Cardiac MRI was performed at 1 week post-MI, before treatment, and at 3 and 6 weeks post-MI, corresponding to 1 and 4 weeks after treatment, respectively. Mice received either an A5:ETE-BuSA drape electrode without mRNA or a VEGF-A mRNA-loaded A5:ETE-BuSA drape electrode. Representative cardiac MRI images are shown for the (a) A5:ETE-BuSA and (b) VEGF-A mRNA-loaded A5:ETE-BuSA group at each time point (Green line: Epicardium, Red line: Endocardium). Longitudinal quantification of (c) ejection fraction, (d) stroke volume, (e) end- diastolic volume and (f) cardiac output at 1, 3 and 6 weeks post-MI. Reference ranges for healthy C57Bl6N mice are given (N = 10). Data are presented as mean ± SD. N = 7 for A5:ETE-BuSA, N = 8 for A5:ETE-BuSA + VEGF-A. Statistical analysis was performed using Repeated Measures ANOVA with LSD post-hoc testing: * denotes p <0.05 between time points 1 and 3 or 6 within the VEGF-A mRNA-loaded A5:ETE-BuSA group; $ denotes p <0.05 between time points 3 and 6 within the VEGF-A mRNA-loaded A5:ETE-BuSA group; & denotes p <0.05 between experimental groups.

### Therapeutic Effects of VEGF-A mRNA delivery via A5:ETE-BuSA in Myocardial Infarction

To determine whether the functional improvements observed by cardiac MRI were accompanied by structural remodeling of the infarcted heart, we examined ventricular morphology and tissue composition at 6 weeks post-MI. The ventricular wall thinning characteristic of post-infarction remodeling where H&E staining revealed pronounced thinning of the ventricular wall in animals receiving the A5:ETE-BuSA drape electrode without VEGF-A mRNA. In contrast, ventricular morphology was better preserved in animals treated with the VEGF-A mRNA-loaded A5:ETE- BuSA drape electrode. (Figure 5a). Regional ventricular wall thickness was subsequently quantified from the cardiac MRI datasets using a previously described multiplanar approach^23^. The LV was assessed across short-axis slices spanning the apical-to-basal axis, with each slice divided into six sectors (Figure 5b). Smooth bulls-eye views of mean end-diastolic wall thickness demonstrated that remodeling was concentrated in the infarct-affected regions (Figure 5b), which were therefore selected for quantitative analysis. VEGF-A mRNA delivery significantly preserved lateral wall thickness compared with A5:ETE-BuSA treatment alone, consistent with the histological findings (Figure 5b). In contrast, septal wall thickness did not differ between groups, in line with the predominantly lateral LV territory affected by LAD ligation^32^. Longitudinal analysis further supported this regional effect. In animals receiving A5:ETE-BuSA alone, lateral wall thickness was significantly lower than septal wall thickness at both 3 and 6 weeks post-MI (Figure S6a). This regional disparity^32^ was absent in animals treated with VEGF-A mRNA (Figure S6b), indicating sustained preservation of the infarct-affected ventricular wall. Moreover, lateral wall thickness positively correlated with ejection fraction across animals (r = 0.5977, P = 0.0145; Table S2). These structural findings provide a mechanistic context for the functional recovery observed by cardiac MRI. Following MI, loss of viable myocardium and progressive ventricular remodeling result in infarct-wall thinning, scar expansion, and impaired contractile function^33^. Here, a single administration of VEGF-A mRNA via the A5:ETE-BuSA drape electrode attenuated regional ventricular thinning, with the strongest effect in the lateral infarct territory. The association between preserved lateral wall thickness and ejection fraction further suggests that attenuation of adverse ventricular remodeling contributed to the improvement in global cardiac function.

**Figure 5.**
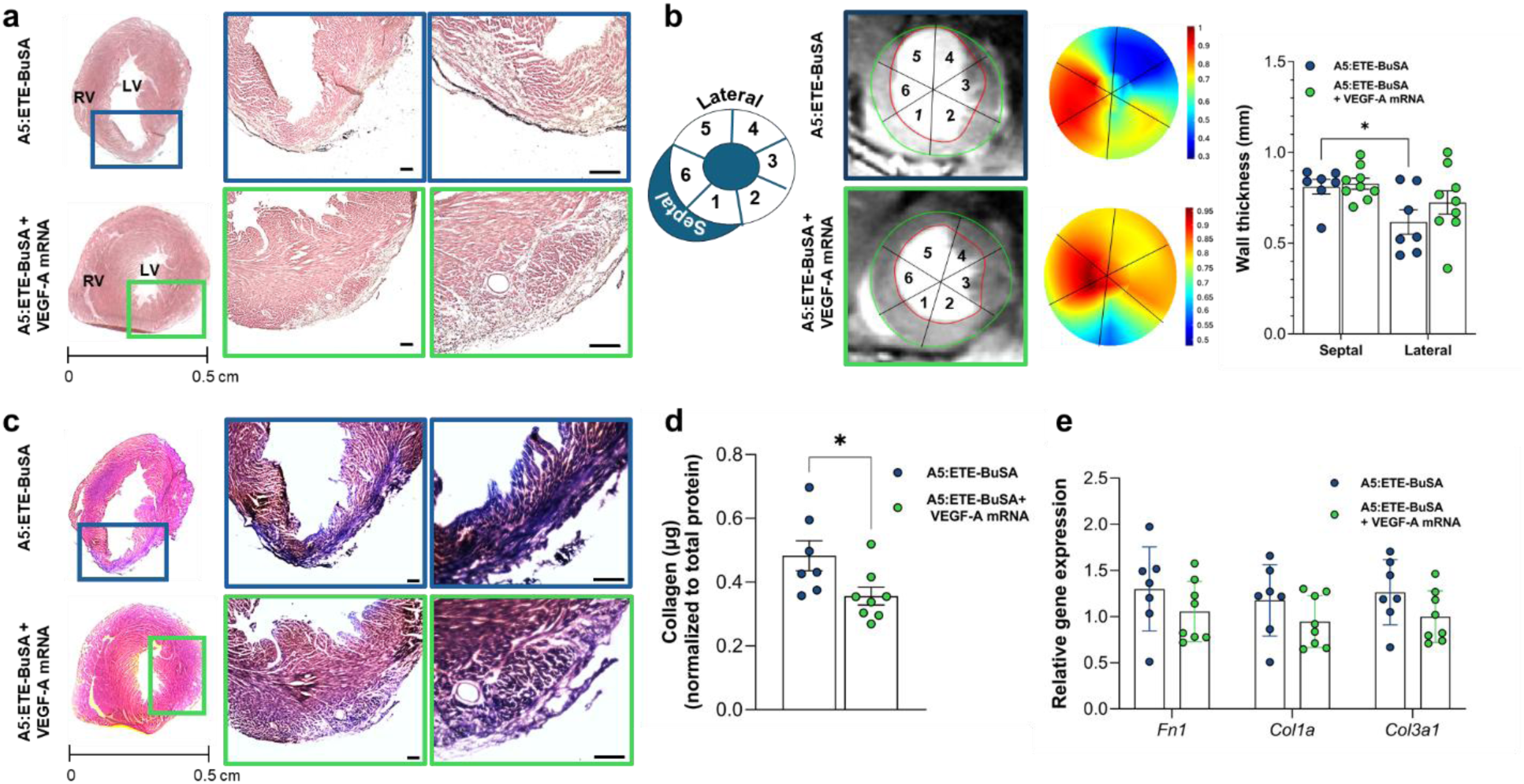
VEGF-A mRNA delivery preserves left ventricular wall thickness and attenuates myocardial fibrosis after MI. (a) Representative hematoxylin and eosin (H&E)-stained heart sections showing ventricular morphology in the indicated experimental groups. (b) Schematic illustrating the short-axis slices and sector-based segmentation used for regional wall-thickness analysis. Representative cardiac MRI images illustrating the sector-based assessment of left ventricular wall thickness and corresponding bull’s-eye plots showing mean end-diastolic wall thickness across the left ventricle. Quantification of end-diastolic wall thickness in septal and lateral sectors across the selected short-axis slices. (c) Representative Masson’s trichrome (MTC)- stained heart sections showing myocardial fibrosis. (d) Quantification of myocardial collagen content using a Sirius Red-based colorimetric assay. (e) Relative expression of fibrosis-associated genes: fibronectin (*Fn1*), Collagen type 1 alpha (*Col1a*) and Collagen type 3 alpha (*Col3a*). Data are presented as mean ± SD. Data are presented as mean ± SD. N = 7 for A5:ETE-BuSA, N = 8 for A5:ETE-BuSA + VEGF-A. Statistical analysis was performed using two-way ANOVA for panel b and a t-test in panels d and e. * denotes p<0.05. Scale bar 200 µm.

Next, we evaluated myocardial fibrosis using Masson’s trichrome (MTC) staining. Hearts from the A5:ETE-BuSA group exhibited extensive fibrosis within the LV (Figure 5c), whereas fibrotic tissue was lower in the VEGF-A mRNA group (Figure 5c). This finding was confirmed by independent quantification of collagen content using a Sirius Red-based colorimetric assay, which demonstrated significantly lower collagen levels in VEGF-A mRNA-loaded A5:ETE-BuSA - treated hearts (Figure 5d). Relative LV expression of fibrosis-associated genes showed a trend toward lower fibronectin (Fn1), Collagen type 1 alpha (Col1a), and Collagen type 3 alpha (Col3a) transcript abundance after VEGF-A mRNA delivery (Figure 5e). Together with the preservation of LV wall thickness, these findings indicate that localized VEGF-A mRNA delivery attenuated adverse post-MI remodeling. Excessive extracellular matrix deposition and collagen accumulation contribute to ventricular stiffening, wall thinning, and impaired contractile function after MI^34^. Thus, the reduction in fibrosis observed here provides a structural correlate for the improved ventricular function detected by cardiac MRI. To further assess cardiac stress, left ventricular NT- proBNP levels as well as *Nppb,* and *Fabp3* expression were measured (Figure S7). All parameters tended to decrease following VEGF-A mRNA treatment, although neither difference reached statistical significance. Across animals, however, circulating NT-proBNP levels were inversely correlated with ejection fraction (r = −0.5354, P = 0.0485; Table S2), consistent with the relationship between increased cardiac stress and impaired ventricular function. We next examined the vascular response to VEGF-A mRNA delivery. Immunofluorescence staining for vimentin and tomato lectin^35^ revealed lectin-positive structures throughout the myocardium, with a greater abundance of lectin-positive areas in VEGF-A mRNA-treated hearts than in the group receiving VEGF mRNA delivery (Figure S8a,b). In parallel, expression of von Willebrand factor (*vWF*), a marker associated with endothelial activation and vascular injury^36^, was significantly reduced following VEGF-A mRNA delivery (Figure S8c). These findings suggest that VEGF-A treatment was associated with changes in the post-infarct vascular compartment and reduced endothelial stress^36^.

Taken together, the functional improvement following VEGF-A mRNA delivery was accompanied by coordinated structural and vascular changes in the infarcted heart, including preservation of the infarct-affected ventricular wall, reduced collagen accumulation, and altered vascular and endothelial responses. These findings are consistent with previous studies showing that appropriately localized VEGF signaling can attenuate adverse post-MI remodeling^37^. This localized delivery strategy also offers a safety advantage over LNP- or viral vector-based mRNA delivery, as naked mRNA that diffuses beyond the injection site is unlikely to transfect distant tissues. By combining transient VEGF-A expression with electrofunctionalization-assisted, spatially localized delivery, the A5:ETE-BuSA platform conferred functional, structural, and vascular benefits following a single-dose administration.

This localized delivery concept, based on an in situ-formed electroactive mRNA-delivery device, addresses a key translational challenge in mRNA therapeutics: the targeted delivery of biologically active cargo to injured tissue. More broadly, it provides a framework for developing targeted nucleic acid therapies as an alternative to LNPs and viral vectors, with particular potential for minimally invasive treatment of myocardial injury without the need for open surgery.

## Acknowledgement

This study was conducted within the Lund University Strategic Research Areas, MultiPark and NanoLund. Equipment within Lund Nano Lab (LNL) was used to enable this research. RO acknowledges funding from the Swedish Cancer Society (24 3629 Pj) and the Swedish Research Council (2023-04965). MH acknowledges the Swedish Science Council grant no. 2025-05838.

## Author contributions

Conceptualization: R.O., M.H., UA. Methodology: R.O., M.H., A.M., U.A., S.S.G. Cell Work: U.A., K.H., U.D. Electrical characterization: U.A., A.S.Y, M.H., S.S., F.E. Flow Cytometry: A.S.Y., U.D., U.A. Chicken embryo experiments: U.A., K.H., UD. Zebrafish experiments: U.A., K.H. Mouse experiments: S.S.G., U.A. Immunofluorescence: U.A., S.S.G., U.D. Histology: S.S.G. Chemistry: R.O., S.S., U.D. Supervision: R.O., M.H., A.M. Funding acquisition: R.O., M.H., A.M. Writing – original draft: U.A., R.O., M.H., S.S.G., and A.M. Writing, review, and editing: All authors reviewed and edited the draft.

## Data, code, and materials availability

All data supporting the findings of this study are available within the article and its supplementary files. Any additional requests for information can be directed to and will be fulfilled by the lead contact. Source data are provided with this paper.

