## Supplementary document for "Bioresorbable, Injectable In Situ-Formed Electroactive mRNA-Delivering Device for Precision Gene Therapy after Myocardial Infarction"

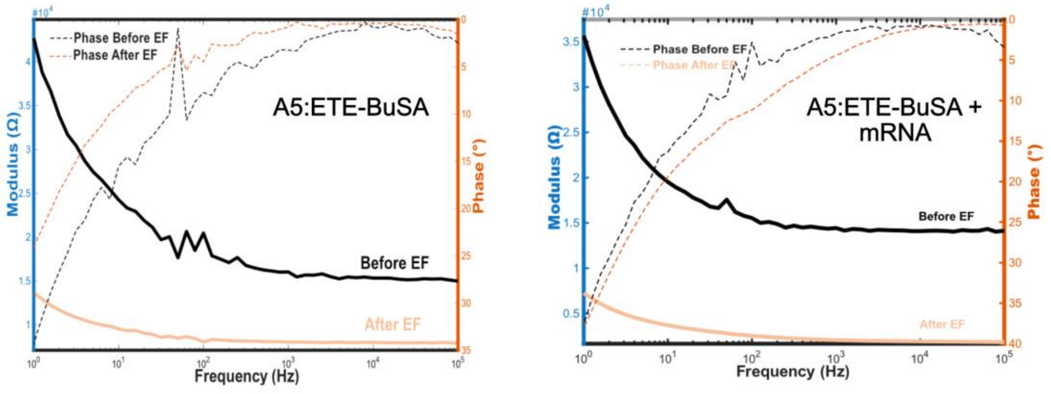


**Figure S1.** Electrochemical impedance spectroscopy (EIS) analysis of the A5:ETE-BuSA and A5:ETE-BuSA+mRNA formulations before and after electrofunctionalization.

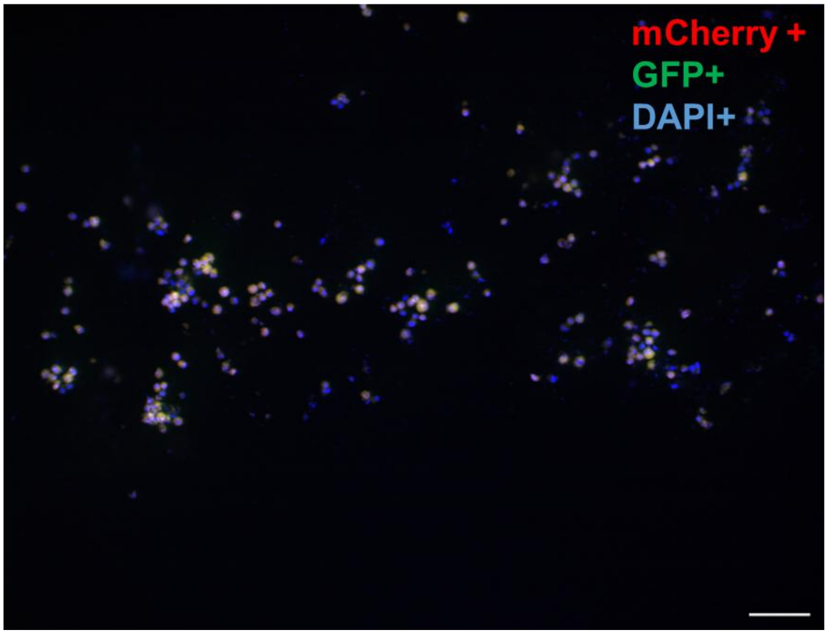


**Figure S2.** Representative fluorescence micrograph showing U87 cells co-expressing GFP and mCherry.

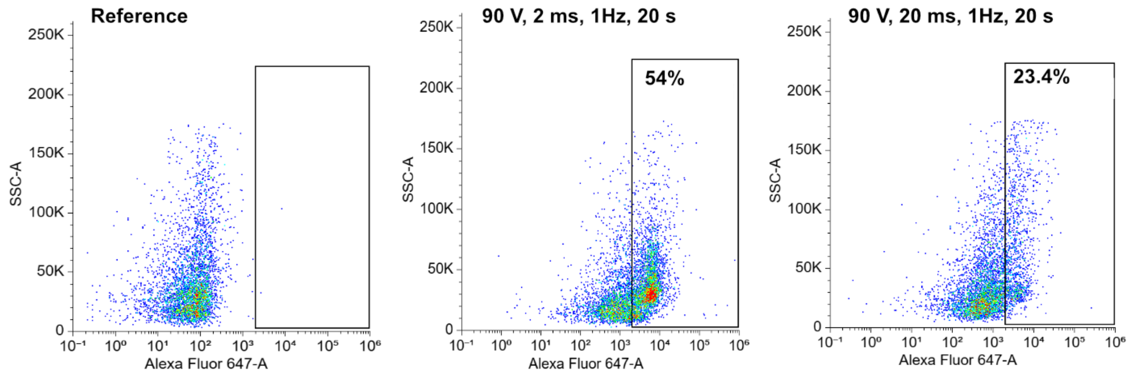


**Figure S3.** Flow cytometry quantification of GFP-positive cells in dissociated CAM tumor preparations after electrotransfection at 90 V.

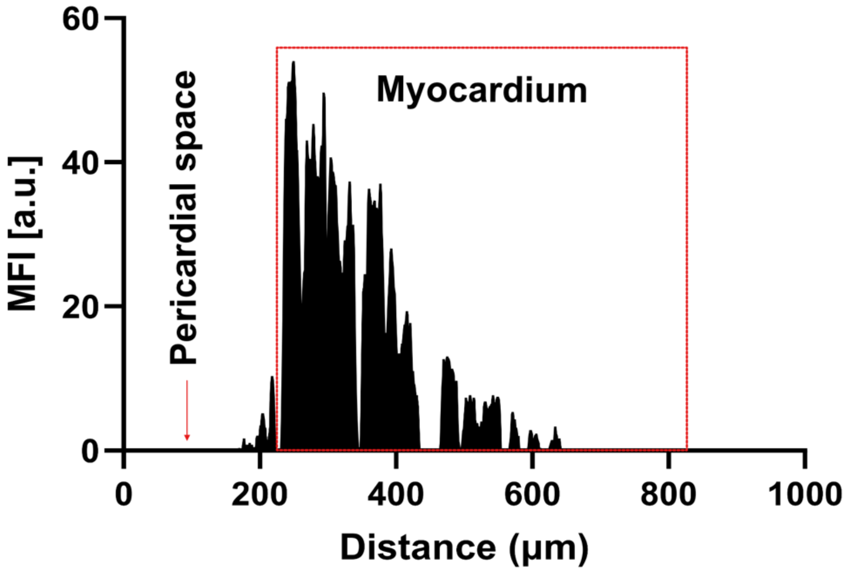


**Figure S4.** Fluorescence intensity (MFI) as a function of distance from the injection site into the porcine myocardium.

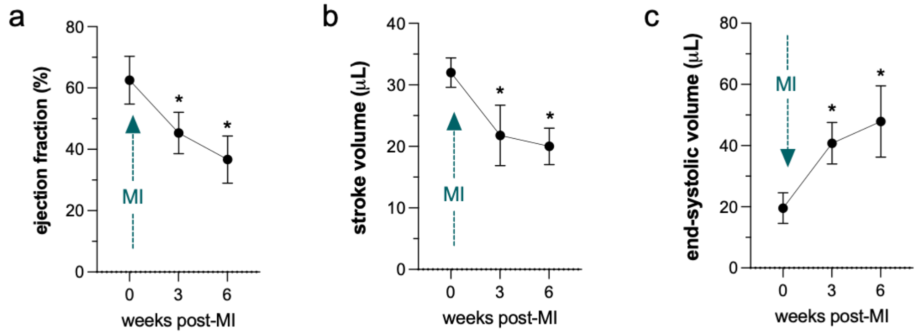


**Figure S5.** Longitudinal assessment of cardiac function pre- and post-myocardial infarction. Cardiac MRI quantification showing pre-and post-MI levels of left ventricular (a) ejection fraction, (b) stroke volume and (c) end-systolic volume with the arrow indicating the time point of myocardial infarction (MI). N = 9. Data are presented as mean ± SD. Statistical analysis was performed using RM-ANOVA with LSD post-hoc testing: * denotes p <0.05 between time point 1 and 3 or 6.

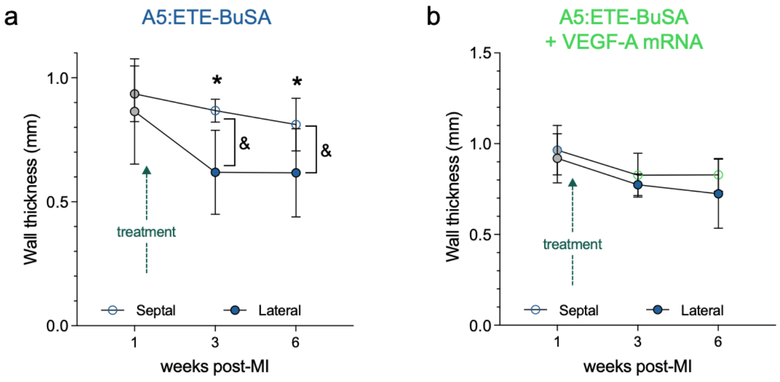


**Figure S6.** Longitudinal regional assessment of ventricular wall thickness following VEGF-A mRNA delivery. Longitudinal assessment of lateral and septal left ventricular wall thickness in animals receiving the (a) A5:ETE-BuSA drape electrode without VEGF-A mRNA and (b) the VEGF-A mRNA-loaded A5:ETE-BuSA drape electrode at the indicated MRI time points. Data are presented as mean ± SD. N = 7 for A5:ETE-BuSA, N = 8 for A5:ETE-BuSA + VEGF-A. Statistical analysis was performed using Two-Way ANOVA with LSD post-hoc testing: * denotes p <0.05 between time point 1 and 3 or 6 within the same region; & denotes p <0.05 between two regions at one time point.

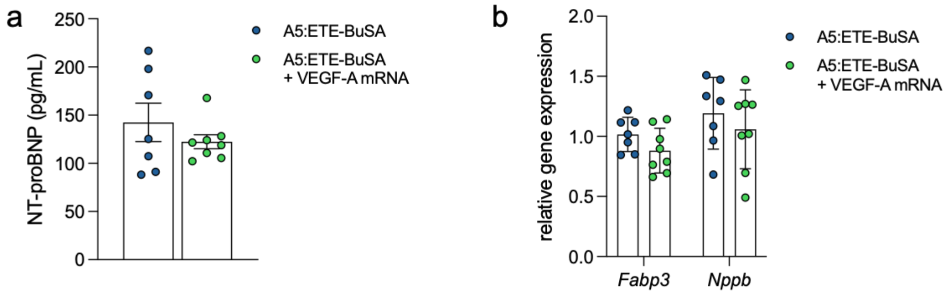


**Figure S7.** Left ventricular NT-proBNP levels following VEGF-A mRNA delivery. **(a)** N-terminal pro-B-type natriuretic peptide (NT-proBNP) levels in left ventricular tissue were quantified by ELISA. **(b)** Relative left ventricular expression of cardiac stress-associated genes: fatty acid binding protein 3 (*Fabp3*) and pro-B-type natriuretic peptide (*Nppb*). Data are presented as mean ± SD. N = 7 for A5:ETE-BuSA and N = 8 for A5:ETE-BuSA + VEGF-A. Statistical analysis was performed using unpaired t-test.

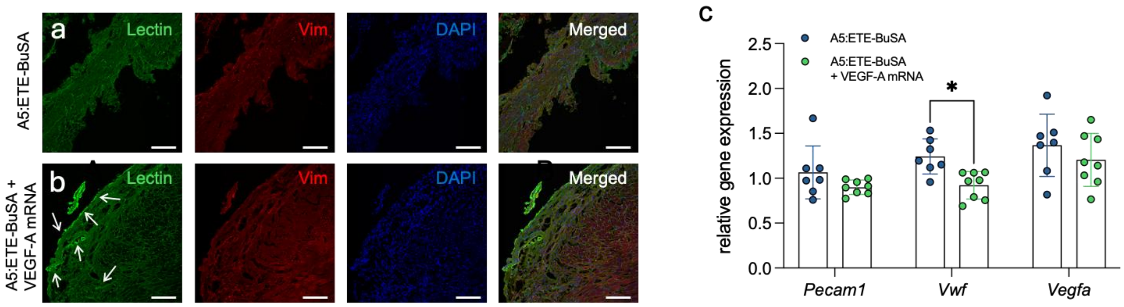


**Figure S8.** VEGF-A mRNA delivery alters vascular features in the left ventricle. (**a** and **b**) Representative immunofluorescence images of heart sections stained for vimentin and tomato lectin, with lectin-positive vascular structures indicated. **(c)** Relative transcript abundance of the vascular-associated genes *Pecam1* (platelet/endothelial cell adhesion molecule 1), *Vwf* (von Willebrand factor), and *Vegfa* (vascular endothelial growth factor A) in left ventricular tissue. Data are presented as mean ± SD. n = 7 for A5:ETE-BuSA and n = 8 for A5:ETE-BuSA + VEGF-A. Statistical analysis for panel B was performed using unpaired t-test. Scale bar is 100 µm

**Supplementary Tables**

**Table S1.** Primer sequences.

| | |
|---|---|
| L14 | F: 5'-GGCTTTAGTGGATGGACCCT-3'<br>R: 5'-ATTGATATCCGCCTTCTCCC-3' |
| Nppb | F: 5'-GAGTCCTTCGGTCTCAAGGC-3'<br>R: 5'-ACTTCAGTGCGTTACAGCCC-3' |
| Fabp3 | F: 5'-AGAGTTCGACGAGGTGACAG-3'<br>R: 5'-TTGTCTCCTGCCCGTTCCAC-3' |
| Fn1 | F: 5'-GGTGTAGCACAACTTCCAATTACG-3'<br>R: 5'-GGAATTTCCGCCTCGAGTCT-3' |
| Col3a1 | F: 5'-CCCTGGACCTCAGGGTATCA-3'<br>R: 5'-GGGTTTCCATCCCTTCCAGG-3' |
| Col1a1 | F: 5'-CCAAGAAGACATCCCTGAAGTCA-3'<br>R: 5'-TGCACGTCATCGCACACA-3' |
| vWF | F: 5'-CTTCTGTACGCCTCAGCTATG-3'<br>R: 5'-GCCGTTGTAATTCCCACACAAG-3' |
| PECAM1 | F: 5'-GCAAATACCCACAGTTCCTCCC-3'<br>R: 5'-GGATGGTGAAGTTGGCTACAGG-3' |
| VEGFa | F: 5'-GGCCTCCGAAACCATGAACT-3'<br>R: 5'-GTCCACCAGGGTCTCAATCG-3' |

**Table S2.** Correlation of cardiac function with circulating NT-proBNP levels and left ventricular wall thickness. Pearson correlation analysis was performed to assess the relationship between ejection fraction (EF) and circulating N-terminal pro-B-type natriuretic peptide (NT-proBNP) levels, as well as between EF and lateral left ventricular wall thickness. Exact correlation coefficients (r) and corresponding P values are reported.

| **Correlation** | **Ejection fraction (%)** | |
|---|---|---|
| | r (Pearson) | P value |
| lateral ventricular wall thickness (mm) | 0.5977 | 0.0145 |
| NT-proBNP (pg/ml) | −0.5354 | 0.0485 |
